# Lactate accumulation induced genioglossus myoblast injury under hypoxia

**DOI:** 10.1101/2023.11.03.565480

**Authors:** Menghan Zhang

## Abstract

Obstructive sleep apnea (OSA) is an increasingly common disorder of repeated upper airway collapse during sleep, leading to chronic intermittent hypoxia and metabolic alteration such as lactate accumulation. Genioglossus (GG), the largest upper airway dilator, is of great significance in maintaining the patency of the upper airway collapse. However, the lactate mechanism in OSA/CIH models is incomplete understood. Here, extracellular and intracellular contents of lactate were detected, and found hypoxia induced lactate accumulation in GG myoblasts. The we treat GG with lactate, and found lactate inhibited myoblasts proliferation and myotubes formation under hypoxia. When the lactate efflux was blocked by monocarboxylic acid transporter 1 (MCT1, a lactate transport) inhibitor AZD3965 (AZD), the expression levels of myogenic markers MyoD, Myogenin and MyHC were reduced. In summary, lactate efflux obstruction and lactate accumulation induced GG myoblast injury under hypoxia. These findings provide a theoretical basis for regulation lactate metabolism to alleviate muscle fatigue in the treatment of OSA.

## Introduction

Obstructive sleep apnea (OSA) is a common sleep disorder characterized by recurrent collapse of upper airway, chronic intermittent hypoxia (CIH), and can lead to hypoxemia and hypercapnia^1-2^. It has been linked to various systemic diseases such as myocardial infarction, congestive heart failure, stroke, and diabetes mellitus^1,3^. The current treatment mainly relieves symptoms and subsequent complication^4^.

Chronic intermittent hypoxia, an important pathological feature of OSA, leads to a series of metabolic abnormalities by controlling the expression of a series of hypoxia response genes^5-6^. Upregulation of glycolysis, inhibition of tricarboxylic acid cycle (mitochondrial oxidative phosphorylation process), accumulation of lactate, compensatory increase in glutamine uptake, etc. Among them, lactate accumulation is an important factor leading to muscle fatigue. Lactate significantly increases and significantly reduces the pH in active muscles, and causing acidosis^7^. On the other hand, lactate can be reabsorbed and utilized by skeletal muscles as a fuel, known as the intercellular lactate shuttle^8^. Lactate can be transported across membranes through a carrier mediated transport pathway^9^. However, the lactate transport mechanism in OSA/CIH models is incomplete understood.

Genioglossus (GG), the largest upper airway dilator, is of great significance in maintaining the patency of the upper airway collapse^10^. In this study, we investigate the lactate metabolism mechanism of GG in CIH mice.

## 2. Material and methods

### 2.1. Isolation and Primary Culture of genioglossus myoblast

C57BL/6 mice male aged 3-4 weeks were anesthetized by pentobarbital sodium. The mice were euthanized using cervical dislocation method, and the skin and cervical membrane of the neck were stripped to expose the genioglossal muscle tissue. Cut the genioglossus muscle to a size of 1 mm3 before rinsing and discarding the supernatant. Add type I collagenase (3 mg/mL) and digest in a 37 °C water bath for 30 minutes, then add 0.05% trypsin and digest again at 37 °C for 30 minutes. Pure myoblasts can be obtained after two differential adhesion attempts.

### 2.2. Myogenic differentiation assay

The mouse myoblast C2C12 was purchased from the Cell Resource Center of Chinese Academy of Sciences, Shanghai Institute of Life Sciences. Myoblast seeded in expansion medium consisting of α-MEM supplemented with 10% FBS for adherence. When the confluence reached 80%, the culture medium were replaced first with myogenic induction medium containing the following for 24 h: IMDM (Gibco, NY, USA) + 2% HS + 1% PS.

### 2.3. Lactic acid content assessment

Lactic acid content was assessed using lactic acid content assay kit in accordance with the manufacturer’s instructions. For cells: the volume of the extraction solution 1 should be in a ratio of 5-10:1 (recommended 5 × Add 1 mL of extraction solution to 106 cells). then the cells were broken by ultrasound on the ice. Centrifuge to collect the supernatant, then slowly add 0.15mL of extraction solution 2, and centrifuge to obtain the supernatant for testing. For liquid: take 100uL of liquid and add 1mL of extraction solution 1. Centrifuge 12000g at 4 °C for 10 minutes. Take 0.8mL of supernatant, then slowly add 0.15mL of extraction solution 2. Slowly blow and mix until no bubbles are formed. Centrifuge 12000g for 10 minutes and take the supernatant for testing.

### 2.4. Cell Counting Kit-8 assessment

Cell were seeded into a 96 hole plate. Preincubate the plate in a damp incubator. Add 10 μL to each hole with CCK8 solution. Be careful not to introduce bubbles in the holes as they can interfere with the O.D. reading. Incubate the culture plate in the incubator for 1-4 hours. Use an enzyme-linked immunosorbent assay to measure the absorbance at 450 nm.

### 2.5. Immunofluorescence

The cells were fixed with 4% cold polyformaldehyde for 20 minutes and wash with PBS three times. then the cells were transparented for 10 minutes and blocked with BSA. Then the cells were incubated with the first antibody at 4 °C overnight. The first antibody: Added in the second-fluorescence at room temperature for 2 hours (in dark).

### 2.6. Quantitative polymerase chain reaction (qPCR)

Total RNA was isolated from cultured cells using Trizol reagent (Invitrogen, Carlsbad, USA). RNA was reverse-transcribed. Resulting cDNA was then diluted 1:20, and amplified with qPCR using SYBR Green (Tiangen Biotech Co., Ltd.). Cycling conditions: Heat ramp 95°C x 10min, extension (95°C x 10s, 60°C x 32s) x 40 cycles. Primers sequences can be found in Table S1. Fold change gene expression was calculated by normalisation to β-actin. The relative mRNA expression was calculated by using 2^-ΔΔCq^ method.

### 2.7. Statistical analysis

All data were present as the mean ± standard deviation, and analyzed using GraphPad Prism 5 (GraphPad Software, Inc.). T test was used to analyze intracellular contents of lactate, while two-way ANOVA with Dunnett multiple comparisons was used to analyze extracellular contents of lactate, CCK8 OD. One-way ANOVA with Dunnett multiple comparisons was used to measure mRNA. *P* < 0.05 was considered a statistically significant difference.

## 3. Results

### 3.1. Hypoxia increase the content of lactate in GG myoblasts

To investigate the change of lactate in GG myoblasts culture under hypoxia, extracellular and intracellular contents of lactate were measured. It is found that hypoxia increased the intracellular contents of lactate in GG myoblasts culture when compared with normoxia (Fig. 1A). The extracellular content of lactate was significantly increased over time, and hypoxia increased lactate content significantly (Fig. 1B).

**Figure 1.**
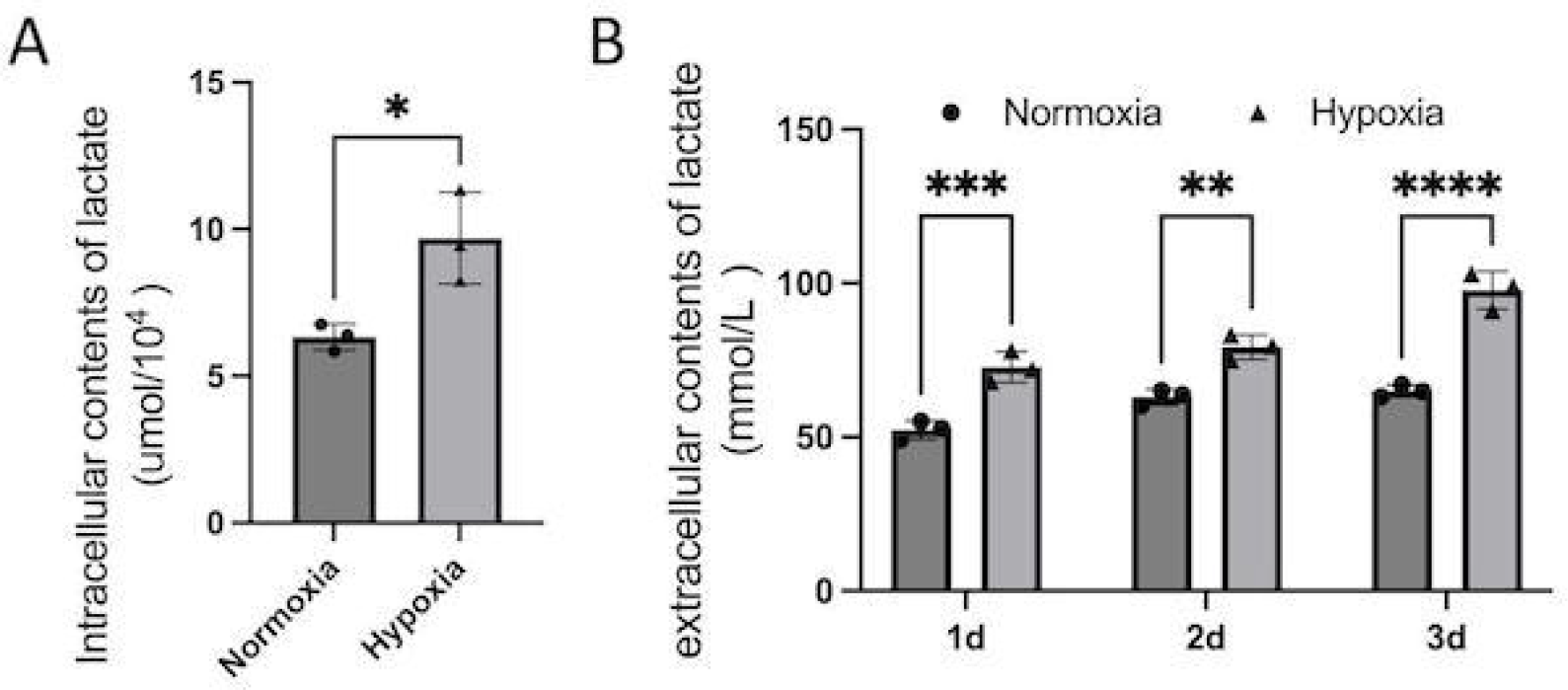
The determination of lactate in genioglossus myoblast under hypoxia. A. Extracellular content of lactate. B. Intracellular content of lactate. \**P* < 0.05, \*\**P* < 0.01 and \*\*\**P* < 0.001.

### 3.2. lactate inhibited myoblasts proliferation under hypoxia

To investigate the effect of lactate on myoblasts proliferation, we examined the effects of different concentrations of lactate on cell proliferation. Can be observed under light microscope that: that hypoxia inhibited cell proliferation, while lactate significantly inhibited myoblast proliferation at a concentration of 60mM (Fig. 2A). CCK8 experiment yielded similar results (Fig. 2B-C).

**Figure 2.**
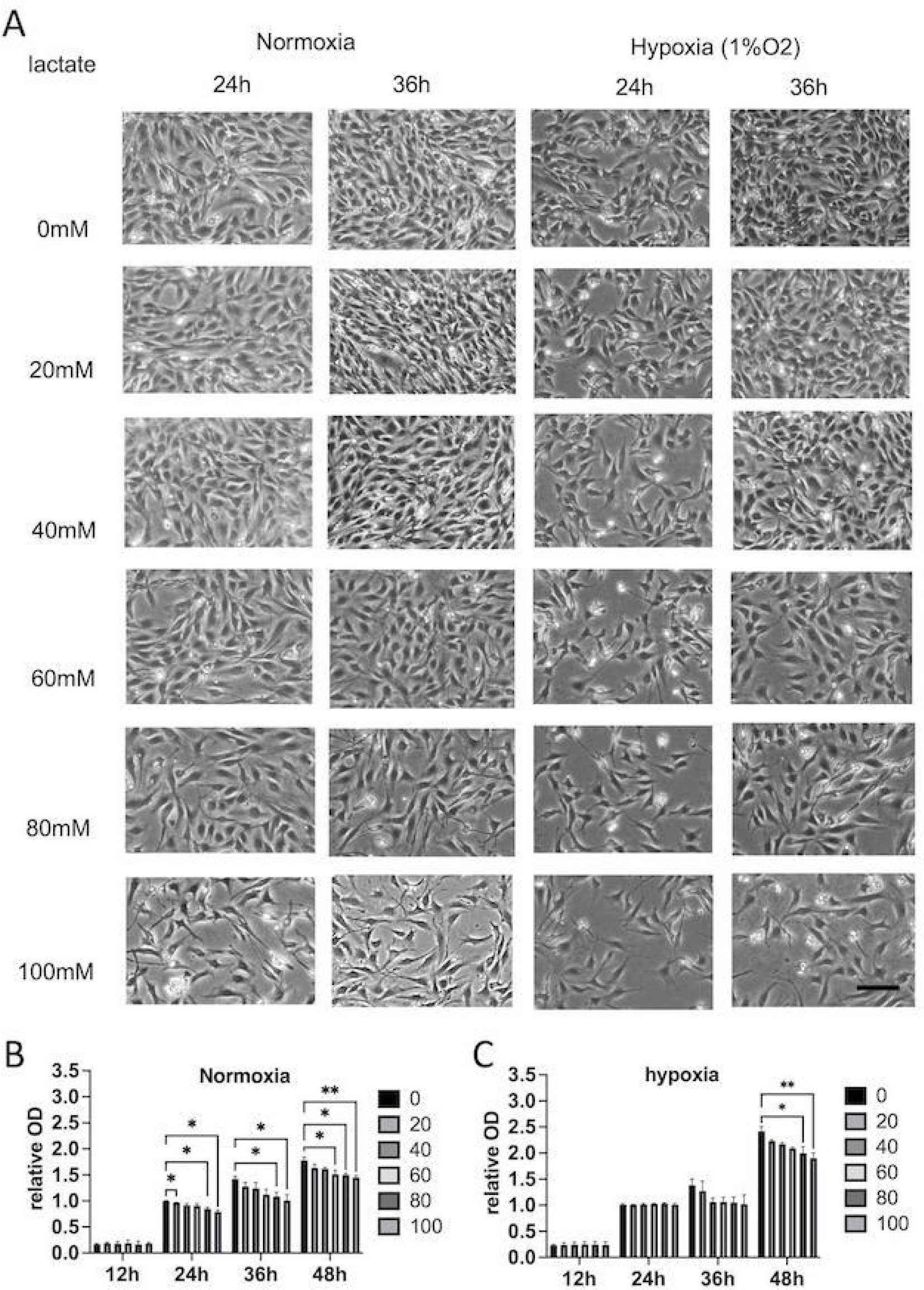
The effect of different concentrations of lactate on cell proliferation. A. Cell proliferation of myoblasts with different concentration of lactate under normoxia or hypoxia cultured 24 h, 48 h and 36 h. B. CCK-8 detects the activity of myoblasts

### 3.3. lactate impeded myotubes formation under hypoxia

To further investigate the effect of lactate on myogenic differentiation, we tested the formation of myotubes (Fig. 3A). Microscopic examination showed that hypoxia had an impact on myotubes formation (Fig. 3B). To quantify the number of myotube formation, immunofluorescence staining of terminal differentiation markers termed myosin heavy chain (MyHC) was performed. The results showed that the number of nuclei fused into MyHC-positive myotubes significantly decreased at the concentration of 20mM and 40Mm lactate, indicating that lactate inhibited myotube formation (Fig. 3C).

**Figure 3.**
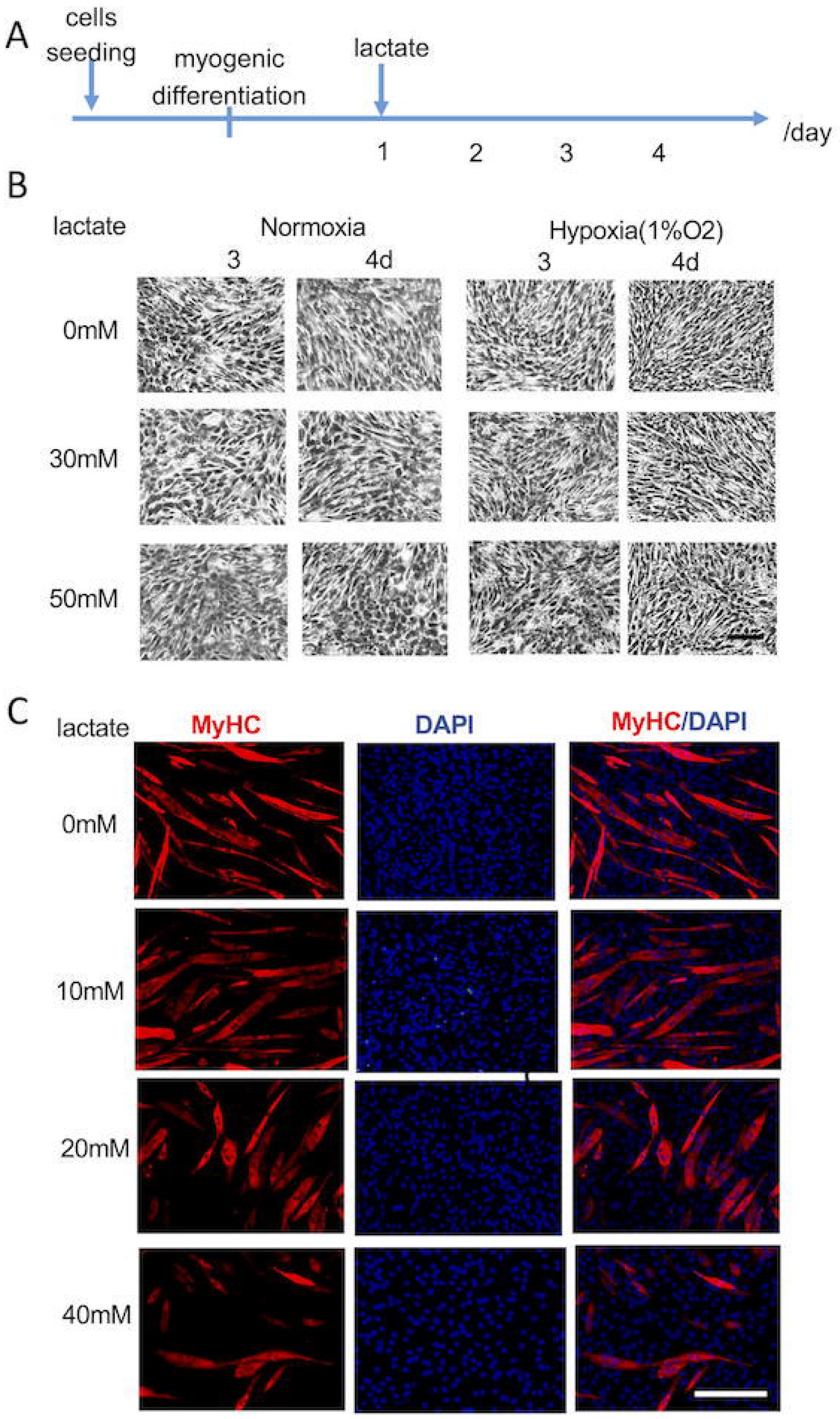
The effect of different concentrations of lactate on myogenic differentiation. A. Schematic diagram of myogenic differentiation and lactate treatment. B. Cell morphology of C2C12 myogenic differentiation by different concentrations of lactate under normoxia or hypoxia cultured 3 d, and 4 h. Scale bar, 200 μm. Scale bar, 200 μm. C. Myotubes and cell nuclei was assessed using immunofluorescence staining with MyHC (red) and DAPI (blue) by different concentrations of lactate. Scale bar, 200 μm.

### 3.4. Inhibiting lactate efflux suppressed myogenic differentiation

Monocarboxylic acid transporter 1 (MCT1) is responsible for lactate transport, by mediating the lactate efflux produced by cellular glycolysis, thus maintain the glycolysis and prevent cell acidification. AZD3965 (AZD) is a MCT1 inhibitor. Therefore, we test the effect of AZD on myogenic differentiation using qPCR of MyoD, Myogenin and MyHC and immunofluorescence staining of myogenin. The results showed that the number of myogenin-positive nuclei decreased at AZD treatment (Fig. 4A). AZD inhibited relative mRNA expression of MyoD (Fig. 4B) and Myogenin (Fig. 4C), indicating that lactate efflux obstruction inhibited myogenic differentiation.

**Figure 4.**
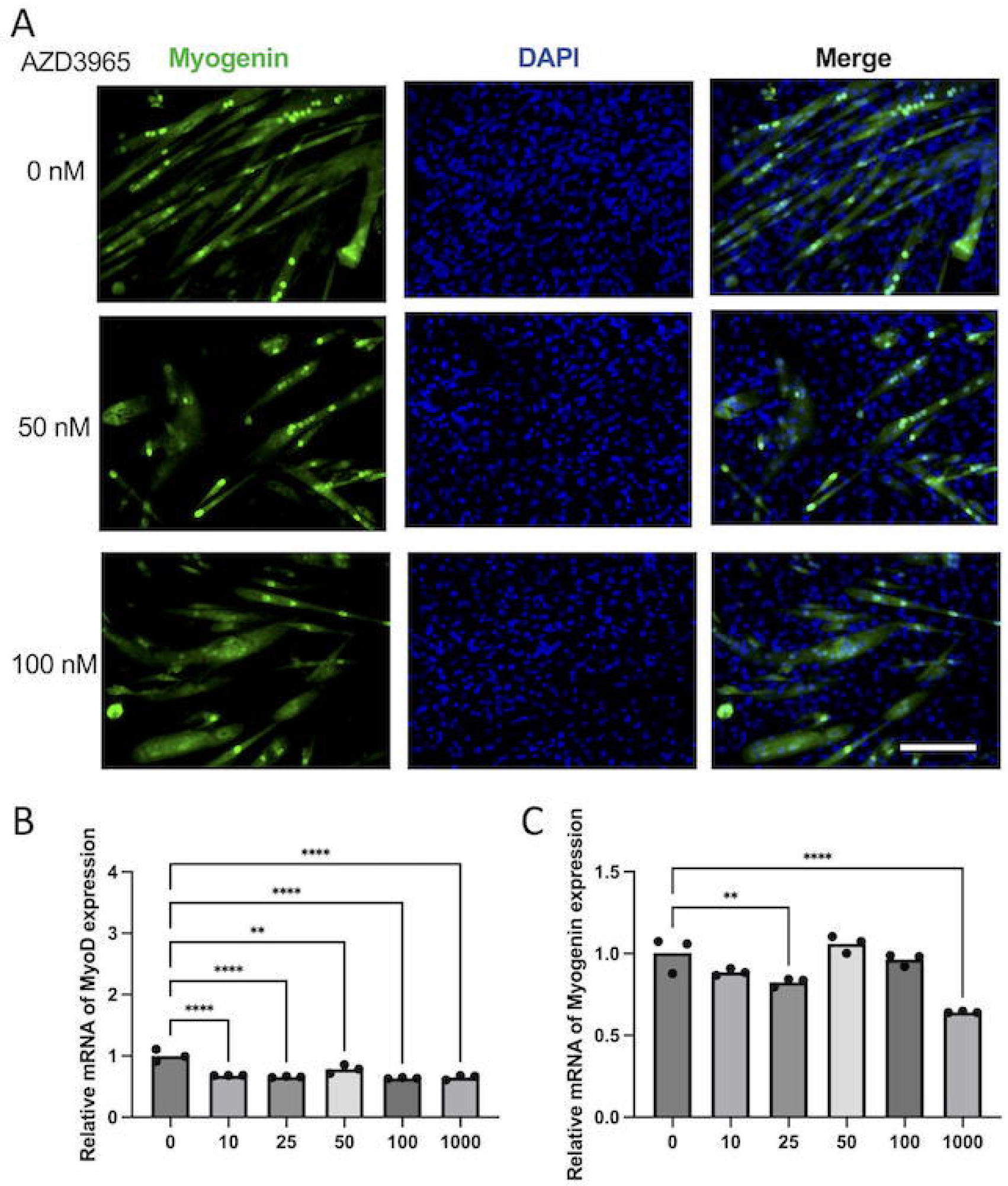
the effect of AZD on myogenic differentiation. A. Immunofluorescence staining with myogenin (red) and DAPI (blue) by different concentrations of AZD. Scale bar, 200 μm. B-D. Relative mRNA expression of MyoD (B), Myogenin (C) and MyHC (D) was assessed using q-PCR. \**P* < 0.05, \*\**P* < 0.01 and \*\*\**P* < 0.001.

## 4. Disscusion

In this study, we found that hypoxia increase the content of lactate in GG myoblasts culture. Furthermore, we found that lactate inhibits myoblasts proliferation and myotubes formation under hypoxia. When the lactate efflux was blocked by MCT1 inhibitor AZD, the myogenic differentiation was inhibited.

Lactate can also have long-term effects on gene expression under hypoxia^11^. Our previous research found that hypoxia increases the expression of HIF-1α^12^. The regulation of HIF-1 under hypoxic conditions promotes the formation and accumulation of LA through increased glucose transport and glycolytic activity. Lactate, as a metabolic product, can also cause hypoxic reactions that are not dependent on HIF-1α. Under hypoxic conditions, accumulated lactate binds and stabilizes NDRG3 protein, thereby activating RAF/ERK signaling mediated lactate triggered hypoxia reactions^13^. In this study, we found that hypoxia leads to an increase in lactate production in GG cells.

Acidosis occurs when acid builds up or when bicarbonate (a base) is lost. The significant increase in lactate caused by metabolic abnormalities can reduce the pH value of active muscle, leading to acidosis^14^. Some studies have shown that acidosis is an important cause of fatigue. The high carbonation environment of OSA patients leads to a low pH state. At the same time, the release of H^+^ from lactic acid can exacerbate this condition^15^. Acidosis is the main cause of fatigue, but there are still many studies opposing this view^16^. Fatigue appearance was in the absence of acidosis. lactic acidosis may actually benefits in delaying fatigue episodes^17^. MCT1 and MCT4 are involved in the proton-dependent transport of monocarboxylates such as L-lactate, which play an essential role in cellular metabolism and pH regulation^18^. In our study, lactate efflux was blocked by MCT1 inhibitor AZD, and the expression of myogenic genes were decreased. This implied that lactate efflux obstruction inhibited myogenic differentiation.

## 5. Summary

Hypoxia induced lactate accumulation in GG myoblasts, and lactate inhibits myoblasts proliferation and myotubes formation under hypoxia. When the lactate efflux was blocked by MCT1 inhibitor AZD, the myogenic differentiation was repressed. In the future, we will further explore the mechanism of lactate transport in the genioglossal muscle tissue of OSA patients, and hope to regulate lactate transport to alleviate muscle fatigue.

