## Supplementary material for "Lactate accumulation induced genioglossus myoblast injury under hypoxia": Table S1

Menghan Zhang^1^

^1^School of Stomatology affiliated to Medical College, Zhejiang University, Hangzhou, China

School of Stomatology affiliated to Medical College, Zhejiang University, Hangzhou, China. Tel: +86-0571-56377243;.

**Table S1. Primers sequences**

| Gene | Forward （5'‑3'） | Reverse （5'‑3'） |
| --- | --- | --- |
| MyoD1 | GGCAGATGCACCACCAGAGT | GTTTGAGCCTGCAGGACACTG |
| Myogenin | GAGACATCCCCCTATTTCTACCA | GCTCAGTCCGCTCATAGCC |
| β-actin | GGCTGTATTCCCCTCCATCG | GTCCCAGTTGGTAACAATGCC |
